# Spatial Transcriptomics Identify T Cell-Driven Mechanisms of Kidney Damage in Immune Checkpoint Inhibitor-Associated Acute Interstitial Nephritis

**DOI:** 10.1101/2025.10.31.685702

**Authors:** Qian Qin, Lennard Ostendorf, Sophia L Wells, Ce Gao, Miles Tran, Firasat M Alikhan, Xiwen Zhang, Api Chewcharat, Robert S Rider, Sean A Prell, Raad B Chowdhury, Astrid Weins, Sujal I Shah, Kavita Mistry, Teresa Bowman, Alexandra-Chloe Villani, Nicole R LeBoeuf, Umut Selamet, Katherine Scovner Ravi, Elad Sharon, David E Leaf, Dennis G Moledina, Meghan E Sise, Iwijn De Vlaminck, Deepak A Rao, Kevin Wei, Shruti Gupta

## Abstract

**Introduction:** Immune checkpoint inhibitor-associated acute interstitial nephritis (ICI-AIN) is the most common finding on histopathology among patients with ICI-associated acute kidney injury (ICI-AKI). Patients with ICI-AIN often have T cell-dominant infiltration of the kidney and high tissue levels of CXCR3 ligands like CXCL9, 10, and 11; however, the mechanisms of inflammation in ICI-AIN are not well-understood.

**Methods:** We applied a sub-cellular spatial transcriptomics platform (Xenium Prime 5K) to compare the cellular composition of kidney biopsy tissue from patients with ICI-AIN with ICI-treated patients with acute tubular necrosis (ICI-ATN).

**Results:** Across 8 kidney biopsy specimens (4 with ICI-AIN, 4 with ICI-ATN), we analyzed 332,000 cells, comprising kidney parenchymal cells and infiltrating immune cells. Using a spatially-aware cellular neighborhood-based classification, we identified cellular niches corresponding to each part of the nephron, in addition to unique fibrotic and inflammatory niches. Gene pathway analysis identified interferon-gamma (IFN-γ)/STAT1 signaling as strongly increased in ICI-AIN compared to ICI-ATN. While all inflammatory niches were overrepresented in ICI-AIN, CD8^+^ T cell infiltration and proinflammatory myeloid cells were the dominant immune niches. Spatial niche crosstalk analysis revealed that CD8^+^ T cell-derived IFN-γ likely induced a proinflammatory program in myeloid cells, with increased production of CXCL9, 10, and 11. Furthermore, IFN-γ signaling in ICI-AIN was associated with reduced oxidative phosphorylation in kidney tubular niches.

**Conclusions:** Spatial transcriptomics reveal novel insights into key differences in the pathophysiology of ICI-AIN versus ICI-ATN. IFN-γ-producing CD8^+^ T cells are likely key drivers of ICI-AIN and should be investigated as future therapeutic targets.

**Translational Statement:** Spatial transcriptomics may provide insight into differences in the cellular and spatial composition of immune checkpoint inhibitor-associated acute interstitial nephritis (ICI-AIN) and ICI-associated acute tubular necrosis (ICI-ATN). Using the novel Xenium 5K platform, we demonstrate that IFN-γ-producing CD8^+^ T cells are central to the pathogenesis of ICI-AIN, and that CD8^+^ T cell-derived IFN-γ likely induces a proinflammatory state in myeloid cells, with increased tissue production of CXCL9, 10, and 11. We then show that a reduction in oxidative phosphorylation and an IFN-γ-driven influx of proinflammatory myeloid cells may further drive kidney damage in ICI-AIN. Many of these pathways represent potential druggable targets, and our findings may therefore inform future therapeutic approaches for ICI-AIN.

## INTRODUCTION

Immune checkpoint inhibitors (ICIs), which target programmed death (PD)-1, PD-ligand 1 (PD-L1), cytotoxic T-lymphocyte associated protein (CTLA)-4, and lymphocyte activation gene (LAG)-3, release the brakes on the immune system and enhance its ability to destroy tumor cells. These drugs, which are now approved to treat >20 cancer types, have dramatically improved outcomes for a number of malignancies.^1–4^ Acute kidney injury (AKI) is common in the setting of ICIs, with real-world studies showing 15-25% of patients treated with ICIs develop AKI.^5^ While acute interstitial nephritis (AIN) is the most common histopathologic lesion seen in ICI-AKI,^6,7^ AKI can also occur due to non-ICI-related causes, including acute tubular necrosis (ATN). Distinguishing ICI-AIN from ICI-ATN is challenging without a kidney biopsy, as there are no clinical features specific to ICI-AIN.^6^ Furthermore, the mechanisms underlying ICI-AIN are not well-understood.

Recent studies have shown that proinflammatory cytokines like tumor necrosis factor (TNF), interferon-gamma (IFN-γ), and IFN-γ-induced chemokines such as chemokine (C-X-C motif) ligand 9, 10, and 11 are elevated in ICI-AIN but not in ICI-treated patients with AKI from other causes.^8–10^ Single-cell transcriptomic analyses of tissue from patients with ICI-colitis^11^ and myocarditis^12^ have each identified an immune cell infiltration dominated by T cells and inflammatory myeloid cells, in addition to organ-specific differences, such as IL-17A-producing T cells in ICI-colitis.

In contrast to ICI-colitis and ICI-myocarditis, our understanding of the immune cell infiltrate, cellular interactions, and damage pathways in ICI-AIN is limited. Given the expanding indications for ICIs, the burden of disease of ICI-AIN is expected to increase.^13,14^ New insights into underlying mechanisms are urgently needed to improve the identification of at-risk patients, enable non-invasive diagnosis of ICI-AIN, and identify new therapeutic targets without inhibiting the anti-tumor response. Here, we utilized Xenium Prime 5K, a novel spatial transcriptomics technology, to compare and contrast the cellular composition of kidney biopsy specimens from patients with ICI-AIN versus ICI-ATN and identify new mechanisms of kidney damage.

## METHODS

### Patient Selection and Tissue Collection

We identified adult patients (≥18 years old) at Brigham and Women’s Hospital (BWH)/Dana-Farber Cancer Institute (DFCI) who had received ICI therapy within 180 days prior to development of AKI (defined as a ≥1.5-fold rise in serum creatinine [SCr] from pre-ICI baseline)^15^ and who had a kidney biopsy performed for clinical purposes between 2021 and 2024. A total of 8 archival formalin-fixed, paraffin-embedded kidney biopsies were obtained (4 with moderate-to-severe AIN and 4 with ATN, as adjudicated by 2 renal pathologists at BWH). All patients had consented to blood and urine collection, as well as archival tissue collection. Spatial transcriptomic analysis of archival tissue was conducted under IRB approved protocol. (#2023P000393). Additionally, all patients had urine proteomics data available (Olink Target 96 Immuno-Oncology Panel and Olink Target 96 inflammation, with 140 unique proteins), as previously described.^8^

This study was approved by the BWH/DFCI (21-685) and Mass General Brigham Institutional Review Boards (#2021P00044 and #2023P000393).

### Xenium Slide Preparation

Formalin-fixed, paraffin-embedded (FFPE) blocks of kidney biopsies were sliced onto Xenium slides according to the “Xenium In Situ for FFPE-Tissue Preparation Guide” (CG000578 Rev C, 10X Genomics)^16^ protocol. The Xenium slides were then prepared following the “Xenium In Situ for FFPE-Deparaffinization and Decrosslinking” protocol (CG000580 Rev D, 10X Genomics). In brief, the slides were incubated at 60°C for 30 minutes and then sequentially immersed in xylene, ethanol, and nuclease-free water to deparaffinize and rehydrate the tissue. The Xenium slides were then immediately incubated in the decrosslinking and permeabilization solution at 80°C for 30 minutes, followed by a wash with Phosphate-buffered saline (PBS).

The Xenium slides were then processed according to the “Xenium Prime In Situ Gene Expression” user guide (CG000760 Rev A, 10X Genomics) for the remaining slide preparation steps. The samples were hybridized with probes from Xenium Prime 5K Human Pan Tissue & Pathways Panel (PN-1000671, 10X Genomics) at 50°C for 18 hours. After hybridization, the slides were washed and incubated with a ligation reaction mix, followed by another wash step and DNA amplification. Cell segmentation staining was then performed, and the slides were treated with an autofluorescence quencher and DAPI before loading into the Xenium instrument.

### Xenium Analyzer Setup and Data Acquisition

Processed Xenium slides were loaded in the Xenium Analyzer and imaged, following the guidelines in the “Xenium Analyzer User Guide (CG000584 Rev F, 10X Genomics).” After scanning, the Xenium slides were removed from the Xenium Analyzer, and processed with post-run haematoxylin and eosin (H&E) staining according to the “Xenium In Situ Gene Expression – Post-Xenium Analyzer H&E Staining” protocol (CG000613 Rev A, 10X Genomics).

### Xenium Data Analysis

We performed both image-based segmentation CellPose^17^ and transcripts-based cell segmentation Baysor (v0.7.1)^18^ with the prior cell labels from the multi-modal staining information. Then, we integrated the Xenium data with single cell RNA-seq reference atlas through Harmony and projected the labels using nearest neighbours, which are further curated with licensed nephrologists. Spatial domain clustering analysis were performed using the natural boundaries-based method Tessera^19^. Niche were further identified using the de novo clustering analysis and validated with the alignment to H&E imaging. The data analysis is described in more detail in the Supplementary Appendix.

### H&E Imaging and Xenium Cell Alignment

Pre– and post-Xenium H&E imaging scans of the biopsied kidney tissue were collected from all 8 patients. To align H&E and Xenium images, original multi-resolution imaging files were first converted to *png* format. Second, affine transformation was applied by using H&E as the query and the Xenium cells as the target by StAlign.^20^ Landmarks were manually labeled on the corners of both image and Xenium cell positions, then the affine transformation was made on the H&E.

### Using Olink Data to Refine Transcriptomics Analyses

Using Olink data, we focused on proteins that we had previously shown to be differentially abundant in ICI-AIN and ICI-ATN. ^8,21^ For proteins that were included in both the Inflammation and Immuno-Oncology panel, we used the results from the Inflammation panel. We performed pseudo-bulk aggregation of the transcripts per kidney biopsy by the AggregateExpression function in Seurat (v5.3.0)^22^ and normalized expression levels to the biopsy area.

## RESULTS

### Study Design and Cohort Description

We analyzed kidney biopsies of 4 patients with ICI-AIN and 4 with ICI-ATN using the Xenium Prime 5K (10X Genomics) spatial transcriptomics assay to measure *in situ* expression of 5101 genes (**Fig. 1A**). Baseline demographics, characteristics of the AKI event and comorbidities of the patients are shown in **Suppl. Table 1.**

**Figure 1.**
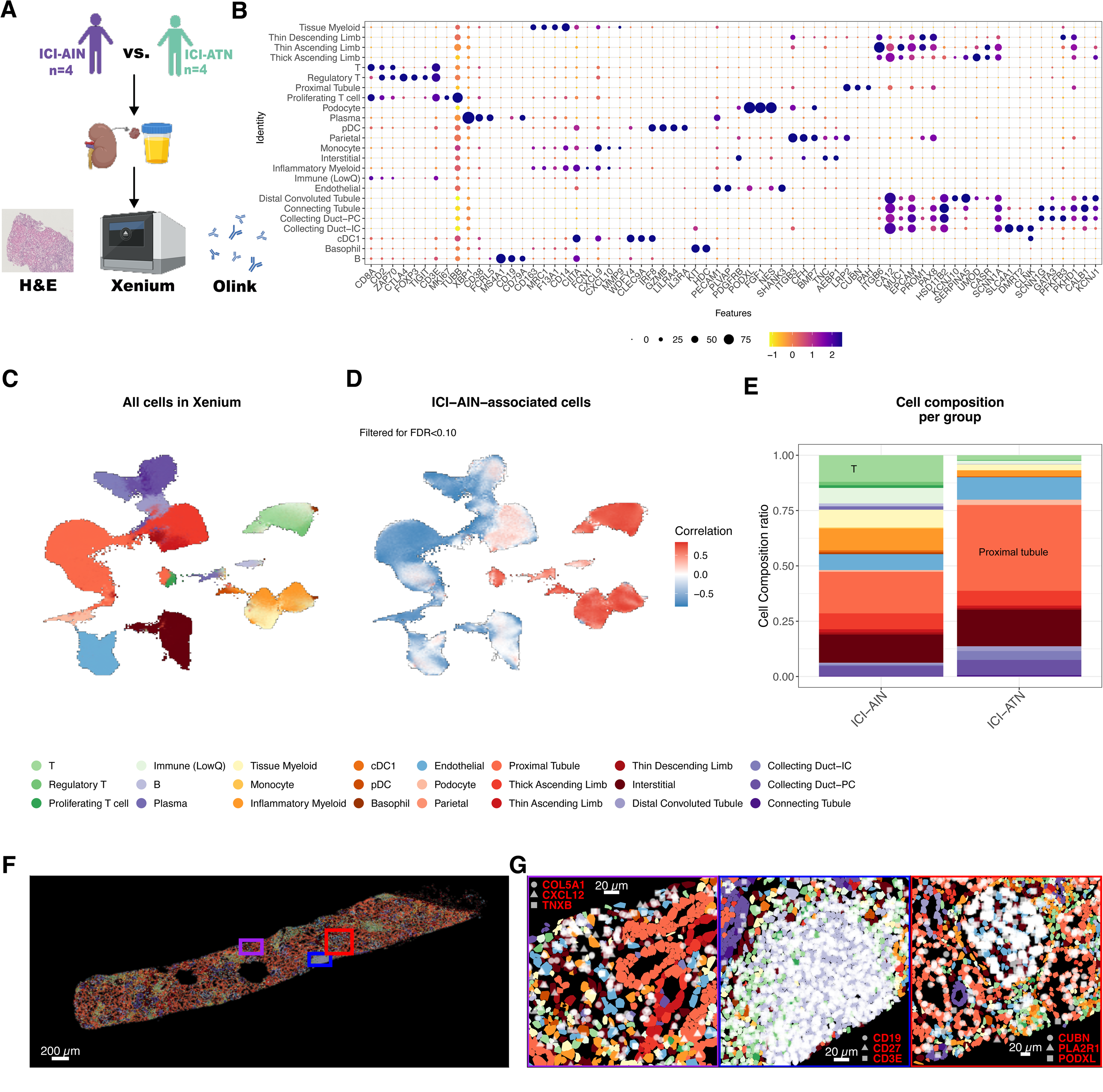
Spatial transcriptomics identifies distinct cell states in ICI-AIN and ICI-ATN. **A.** Study design and techniques involved, **B.** Marker heatmap for the major cell types in spatial transcriptomic data. Rows show the cell types, columns show top selected markers, the size **C.** UMAP of spatial transcriptomic cells with cell type annotation in ICI-AIN and ICI-ATN, the size of the circle indicates the proportion of cells expressing the marker, the color indicates the average expression scale, **D.** Differential abundance of cellular composition measured by ratio over all cells for both groups, **E.** Cell type ratio in all patients of ICI-AIN and ICI-ATN, with corresponding color legend below**. F.** Xenium cells from a representative patient with ICI-AIN, **G.** Zoomed-in regions from F; multiple polygons show the segmented cells in their spatial position; colors of the polygons show the cell type; white dots with different shapes show the transcript positions. Abbreviations: ICI-AIN: Immune checkpoint inhibitor (ICI) therapy-related acute interstitial nephritis; ICI-ATN, Immune checkpoint inhibitor (ICI) therapy-related acute tubular necrosis; pDC: Plasmacytoid dendritic cell, cDC1: conventional dendritic cell type 1, Collecting Duct-PC (principal cells), Collecting Duct-IC (intercalated cell), Immune (LowQ): Immune cell lineage cells with low quality data.

### Immune Cells in ICI-AIN vs ICI-ATN

After cell segmentation and quality control (**Suppl Fig. 1A-D**, **Methods**), followed by label transfer and manual curation strategy (**Methods**), we identified a total of 332,341 cells with 21,251 to 66,751 cells per patient sample. Using the curated cell types in the KPMP atlas as a reference, we defined 24 major cell types, comprising both kidney parenchymal cells and immune cells across all 8 patient samples (**Fig. 1B-C, Suppl. Fig. 1E-F**, **Suppl Fig. 2**).

Next, we applied covarying neighbourhood analysis (CNA)^23^ to identify cell states strongly associated with ICI-AIN and found immune cells to be significantly expanded in ICI-AIN (**Fig. 1D**). Compared to ICI-ATN, ICI-AIN biopsies contained 27-fold more *XBP*^high^plasma cells in ICI-AIN, 8-fold more *IL3RA*^+^ plasmacytoid dendritic cells (pDCs), 7.9-fold more proliferating T cells, 7.6-fold more *FOXP3*^+^ CTLA4^+^ regulatory T cells (Tregs), and 3.6-fold more *CXCL9*^+^*CIITA^+^*inflammatory myeloid cells (**Fig 1E, Suppl. Fig. 1H**, FDR < 0.05 for each).

We also identified more kidney parenchymal cells in ICI-AIN, including *SLC4A1*^+^ Collecting Duct-Intercalated cells (7.7-fold higher), *SCNN1G^+^* Collecting Duct-Principal cells (1.5-fold), *PODXL*^+^ podocytes (3.4-fold), *KCNJ10*^+^ distal convoluted tubule (2.3-fold), and *CUBN*^+^ proximal tubule (2-fold; **Fig 1E, Suppl. Fig. 1E, 1H**). By mapping the cell types and individual detected marker transcripts in space, we were able to confirm the identified cell types in their spatial context, including high levels of *CD19* and *CD3E* transcripts in a large lymphoid aggregate, *PODXL* transcripts in glomeruli, and *CUBN* in proximal tubular cells (**Fig 1F, G**). In summary, we were able to identify characterize the immune cell infiltrate in ICI-AIN using spatial transcriptomics and identified a high density of plasma cells, pDCs, and T cells, among others.

### Immune and fibrosis niches in ICI-AIN vs ICI-ATN

To dissect cellular neighborhoods in their spatial context, we identified cellular niches using the Tessera package.^19^ Tessera is an automated tissue-agnostic tissue segmentation method that introduces novel concepts of spatial cellular neighborhoods or “niches,” delineates the tissues by closed multi-polygons that aggregates spatially coherent cells, and approaches that infer the tissues’ edge with natural tissue boundaries. Tessera enables reducing the sparsity of spatial transcriptomic data, and circumvents potential problems associated with cell segmentation methods by aggregating cells, thereby increasing the number of relevant transcripts per unit and decreasing misattribution errors. Using Tessera to characterize cellular niches in these biopsies, our approach enables us to identify potentially novel cellular niches and biologically meaningful architectural units within the spatial transcriptomic data. In total, we aggregated cells to 22,690 niches. Based on the clustering of these niches based on transcriptional similarities, we defined 9 major spatial niches, including immune, vascular and fibrotic niches, and 6 kidney tissue-related niches (**Fig. 2A**).

**Figure 2.**
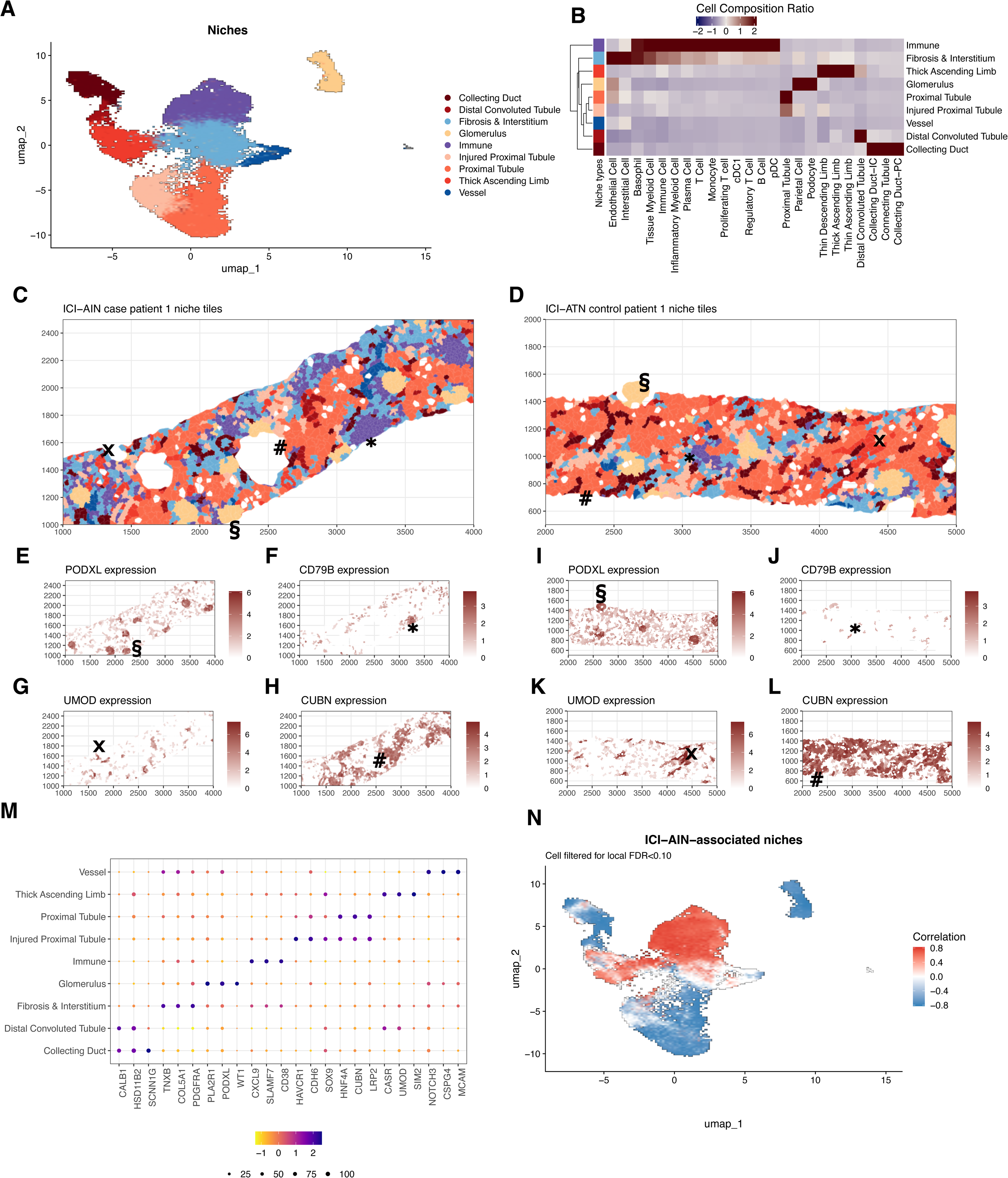
Immune and stromal cellular niches are expanded in ICI-AIN. **A.** UMAP projection of niches using the aggregated gene expression; each dot color represents the annotation of niche type, **B.** Heatmap of cell type ratio in each spatial niche, annotated color bar legend is the same as in A, the cell colors show the abundance of the cell type ratio, **C.** multi-polygon plot visualizes the spatial niches of one representative ICI-AIN patient, **D.** same as C but for a patient with ICI-ATN, **E-L.** marker expression in spatial niches for the same patient in C and D. **M.** CNA of spatial niches, red color shows positively associated niches in ICI-AIN, blue means the negatively associated niches, **N.** Dot heatmap plot showing the marker average expression and expressed tile percentage.

The kidney-related niches included all major parts of the nephron: for example, glomerular niches that were marked by high expression of *PODXL* (a transcript typically found in podocytes) and *WT1* (a marker of parietal epithelial cells), illustrating the multicellular composition in biological units identified by the algorithm (**Fig. 2B-M**). Similarly, we identified niches corresponding to the proximal tubule (with high expression of *CUBN* and *LRP2*), thick ascending limb (marked by *UMOD* and *CASR* expression), distal convoluted tubule (marked by *TRPM6* expression), and collecting duct (expressing *SCNN1G* and *HSD11B2* and combining both intercalated and principal cells). Additionally, we identified niches associated with pathology: injured proximal tubular regions were marked by the expression of markers like *HAVCR1* (encoding Kidney Injury Molecule 1 (KIM-1)) and *SOX9* in addition to the proximal tubule transcriptional signature, as has been extensively studied in the context of AKI.^24–27^ Immune niches contained different immune cells subsets; generally, we identified high levels of IFN-y induced transcripts like *SLAMF7*, *CD38* and *CXCL9*. Similarly, areas of fibrosis were marked by high expression of genes like *PDGFRA* and *COL5A1*.

The identity of the niches was confirmed by their cellular composition (**Fig. 2B**), the spatial expression of key marker genes following the niche distribution (**Fig. 2C-L**), and the comparison of niche distribution and the corresponding H&E slides (**Suppl. Figure 3**).

Applying CNA to characterize cellular niches significantly expanded in ICI-AIN compared to ICI-ATN, we observed an expansion of immune fibrotic niches, as well as niches containing thick ascending limb transcriptional profile in ICI-AIN, while the remaining kidney-specific niches were decreased in ICI-AIN (**Fig. 2N**). The injured proximal tubule niche was more highly represented in ICI-ATN samples. Using a niche-based approach, we were thus able to identify all major parts of the nephron in addition to immune cell and fibrotic niches.

### Immune Niche Subtyping Identifies Key Immunological Processes in ICI-AIN

To better understand the immune cell infiltration in ICI-AIN, we performed unbiased subclustering of the immune niches (**Fig. 3A**). Some niches corresponded to the microenvironment of specific immune cells, such as plasma cell niches with high levels of transcripts as *FCRL5* and *TNFRSF17* (encoding BCMA) or plasmacytoid dendritic cells (expressing *LIL4RA*, and *IL3RA,* and encoding CD123) (**Fig. 3B**). The TLS-like niches showed high expression of genes that are involved in the interaction of B and T cells, such as *CD28* and *ICOS*, as well as their component cells. Other niches showed stronger transcriptional signals of the infiltrated kidney tissue or fibrotic processes in addition to an immune cell signature (termed “CD8 infiltrated kidney” and “inflammation with fibrosis”). We detected two separate types of myeloid cell niches, one with higher expression of pro-inflammatory genes (like *CSF2RB* and *MMP9*) that are often associated with monocyte-derived cells, which we termed “myeloid inflammation,” and another myeloid niche type with increased expression of transcripts associated with tissue-resident or regulatory myeloid cells (such as *CD163*, *MRC1* [encoding CD206] and *F13A1*; M2-like). Notably, T cell-related transcripts such as *CD3E* and *CD8A* were detected in all immune niches to varying degrees, highlighting the role of T cell infiltration as a general feature of ICI-AIN.

**Figure 3:**
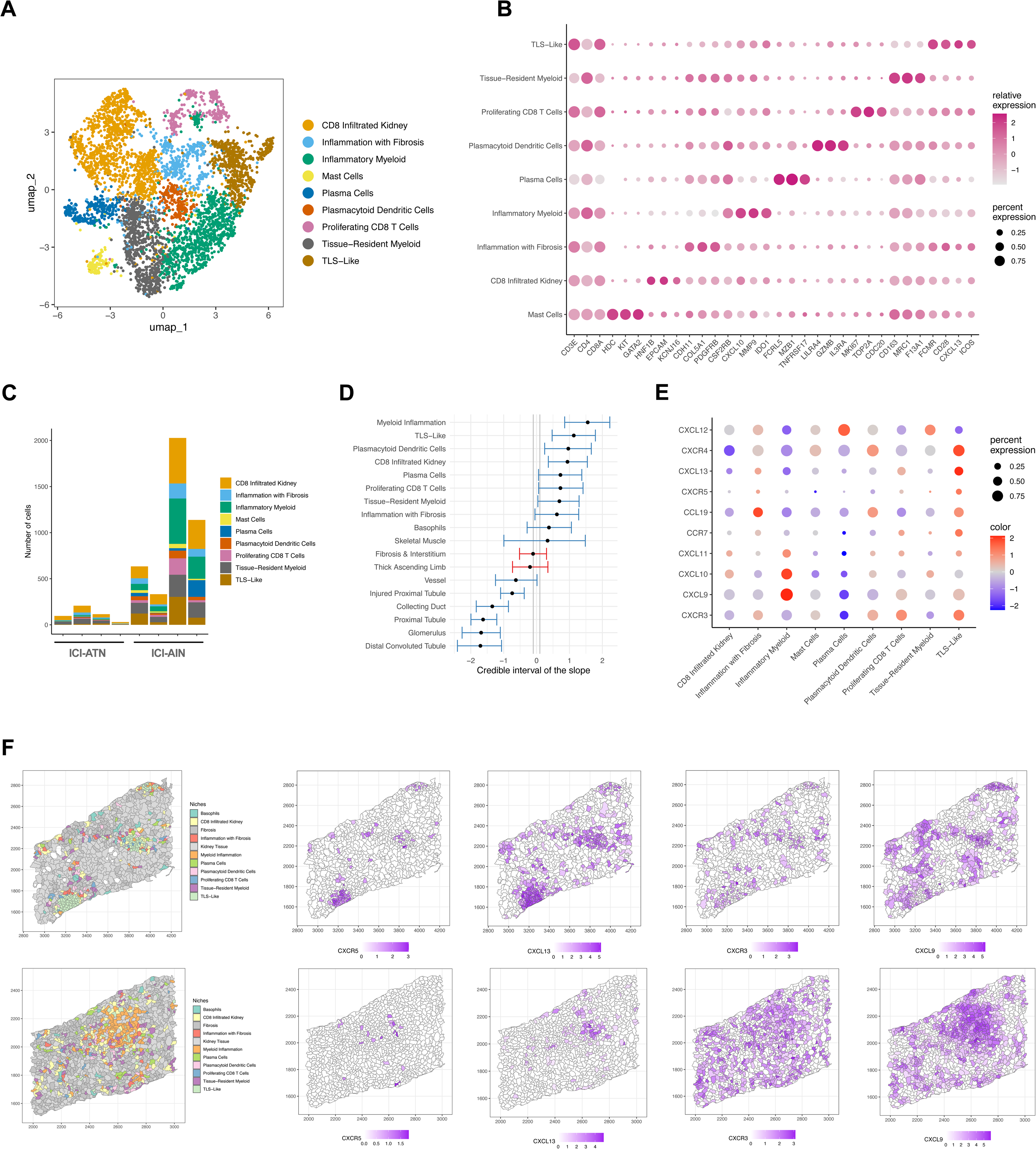
Distinct Immune niche subtypes in ICI-AIN. **A.** UMAP projection of subclusters in ICI-AIN, **B.** Marker genes of immune cell subclusters, **C.** Absolute abundance of immune cell type per biopsy, **D.** Covariate estimates of the sccomp model of niche composition. A positive slope value indicates a positive association with ICI-AIN. Blue color indicates variates passing the significance threshold (*P* < 0.05) **E.** Dot plot of selected chemokine receptor/ligand pairs expressed in immune subniches. **F.** spatial projection of immune subniches and the CXCR5/CXCL13 and CXCR3/CXCL9 chemokine axes.

Compared to ICI-ATN, all immune niche subtypes were increased in ICI-AIN (**Fig. 3C**). In a linear regression model examining the association of specific immune niches with ICI-AIN, all immune niche types were increased in ICI-AIN, with the strongest associations observed for the niche types “inflammatory myeloid”, “TLS-like” niches, and “CD8 infiltrated kidney” (**Fig. 3D**).

### Distinct Chemokine Axes Underlie Immune Niche Subtype Heterogeneity

To better understand drivers of immune niche heterogeneity, we performed ligand-receptor analysis and observed greater overall ligand receptor interaction strength across ICI-AIN vs ICI-ATN (**Suppl. Fig. 4A-B).** In both ICI-AIN and ICI-ATN, CXCL12-CCR4 and CXCL9-CXCR3 interactions were observed in all patients, although the latter pair had higher strengths of association in ICI-AIN compared to ICI-ATN (**Suppl. Fig. 4C-F**). CXCL12-CCR4 was globally present in areas with myeloid inflammation, TLS, and plasma cells in ICI-AIN, whereas it was limited to a few niches in ICI-ATN.

Investigating the different immune subniches, expression of CXCR3 ligands CXCL9, 10, and 11 was found in all subniches, but most prominently in the “myeloid inflammation” niches. were found in all subniches, but most prominently in the “myeloid inflammation” niches. This was associated with massive expansion of CXCR3-expressing cells (**Fig. 3E-F, Suppl. Fig. 4G-J**). The chemokine axes CCR7/CCL19 and CXCR5/CXCL13, both associated with B-T cell interaction and tertiary lymphoid structures, were most strongly detected in the TLS-like immune niches. CXCL12 was most strongly increased in plasma cell niches, as has previously been reported in bone marrow plasma cells.^28^ Our analyses highlight the role of chemokines in the spatial organization of immune infiltrates in ICI-AIN, with CXCR3/CXCL9 interactions in areas of myeloid inflammation and CXCR5/CXCL13 interactions in tertiary lymphoid structures.

### Spatial Transcriptomics Builds on Findings from Urine Proteomics

Building upon our previous urine proteomics analysis,^21^ we compared paired aggregate gene expression and proteomic abundance of 5 differentially abundant proteins in ICI-AIN (CXCL9, CXCL10, CXCL11, FASLG, GZMA) and 1 in ICI-ATN (EGF) (**Suppl. Fig. 5**).^21^

Mean aggregate transcript levels in ICI-AIN for these genes were changed in the same direction as the urinary protein abundance. Gene expression of *CXCL9, CXCL10, CXCL11,* and *GZMA* each completely separated the groups, often showing better separation than urine protein levels.

### Gene Set Enrichment Analysis Identifies Interferon-Driven Responses in ICI-AIN

We next asked what signaling pathways contribute to the immune cell activation and tissue inflammation in ICI-AIN. Comparing differentially expressed genes within the niche types separately, IFN-y-induced genes, such as *STAT1, CXCL9, CD38* and *SLAMF7,* showed the strongest expression in ICI-AIN compared to ICI-ATN in all cellular niches (**Fig. 4A-C, Suppl. Fig. 6-7**). Gene set enrichment analysis of immune-related pathways in the KEGG and Hallmark gene set collections similarly identified an increase in IFN-driven pathways (such “Interferon gamma response,” “chemokine signaling,” “JAK-STAT,” and “antigen processing and presentation”) in ICI-AIN (**Fig. 4A-C)**. Especially within the tubular niches of the kidney, we observed a strong reduction in the expression of genes related to oxidative phosphorylation in ICI-AIN. The spatial expression changes of these signaling pathways were also compared with the aggregated signature expression across the niches (**Fig. 4D-G**). In summary, the IFN-y response signature was the dominant transcriptional immune signature in ICI-AIN.

**Figure 4.**
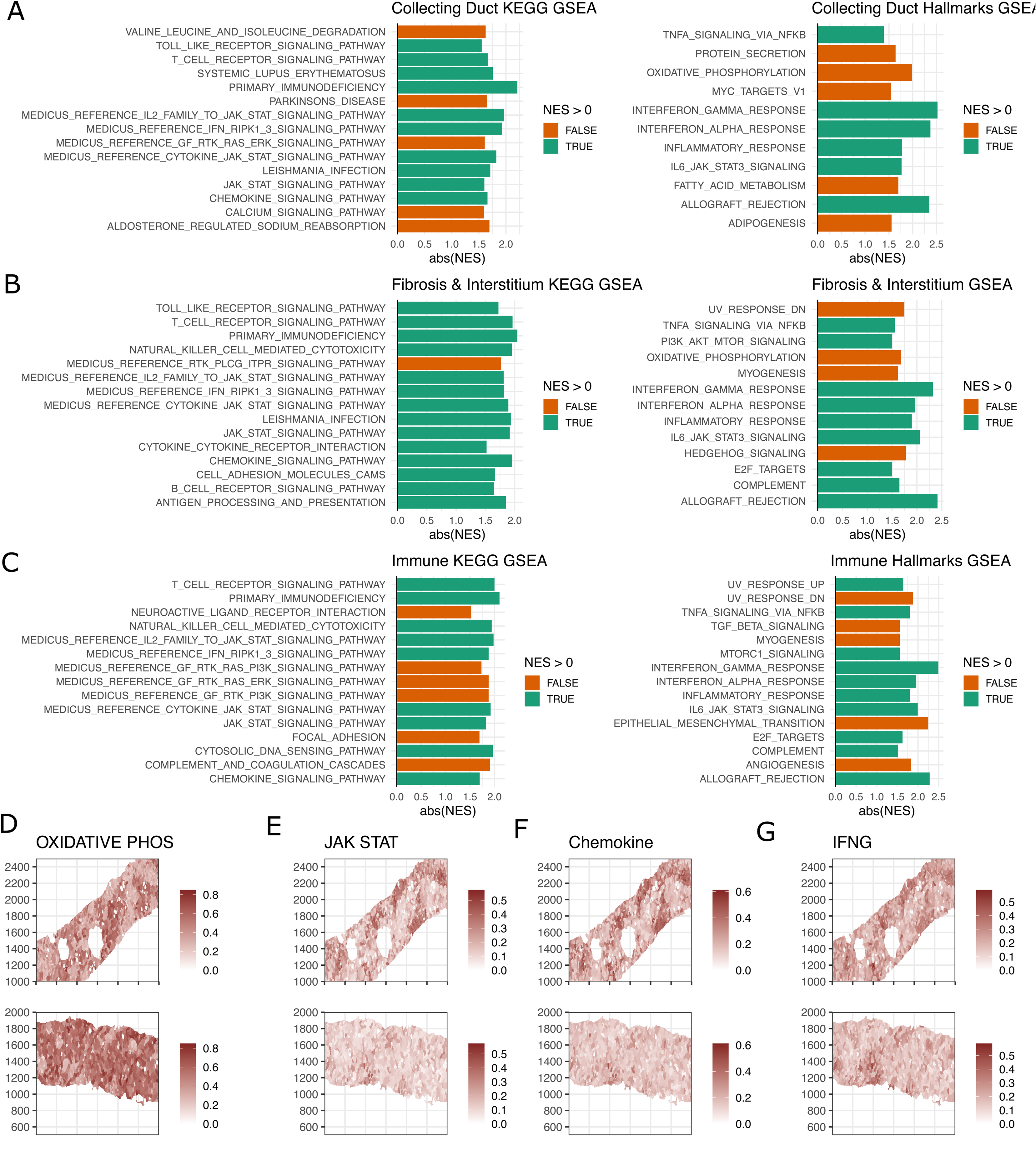
Pathway enrichment analysis based on niche marker expression changes comparing ICI-AIN to ICI-ATN. **A-C**. Left panel is the volcano plot showing the differentially expressed markers in ICI-AIN vs ICI-ATN, the middle and right panels are the KEGG pathway and hallmark enrichment analysis using the ranked-based method GSEA**, D-G,** UCell scores for signatures expression from the top prioritized signalling pathways in representative ICI-AIN (top row) and ICI-ATN (bottom row) samples.

### Tissue-Resident CD8^+^ T Cells in ICI-AIN Produce IFNγ and CXCL13

As we identified IFNγ/STAT1 as the predominant signaling pathway and IFNγ-induced genes as the top DEGs in ICI-AIN, we next investigated the cellular source of IFNγ production. The greatest number of *IFNG* transcripts (75.1% of all *IFNG* transcripts identified in immune cells) were identified in CD8^+^ T cells as compared to other immune cell subsets, including CD4 T cells (**Fig. 5A**). Investigating differentially expressed genes within the CD8^+^ T cells, we found *IFNG* to be one of the top upregulated genes in ICI-AIN (**Fig. 5B**). *CXCL13* was also increased, as were markers of repeated activation such as *PDCD1, LAG3,* and *CTLA4* in ICI-AIN CD8^+^ T cells. While ICI-ATN CD8^+^ T cells expressed markers of short-lived effector T cells like *KLRG1* and *CX3CR1*, CD8^+^ T cells in ICI-AIN showed higher levels of residency program markers like *ZNF683* (encoding HOBIT), *CXCR6,* and *ITGAE* (encoding CD103). Additionally, CD8+ T cells from ICI-AIN patients were enriched for IFN-induced genes like *STAT1*, *GBP1,* and *GBP5*.

**Figure 5.**
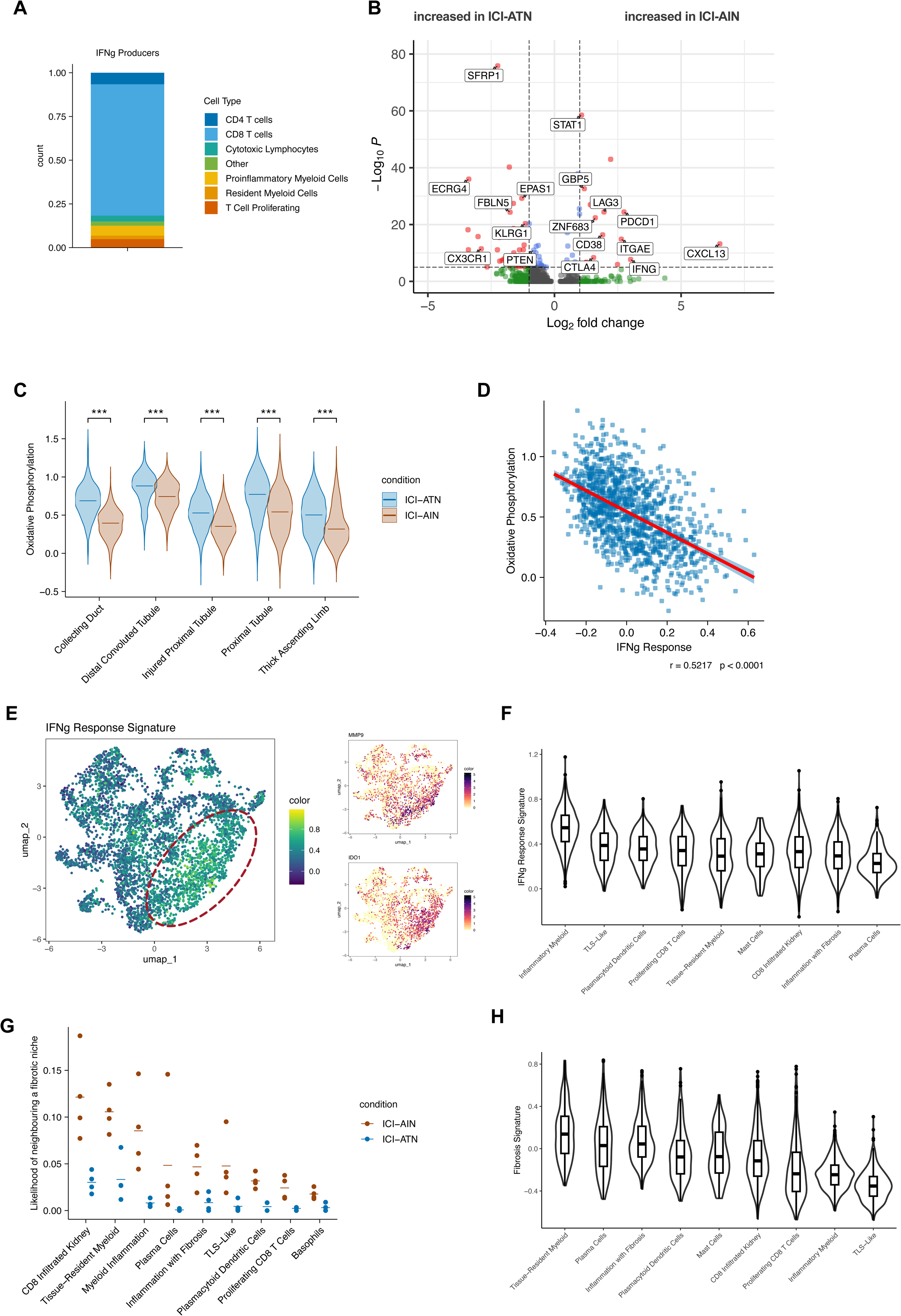
IFNG analysis in the immune and fibrotic niche. **A.** Percentage of different immune cell subsets among all immune cells with *IFNG* transcripts detectable, **B.** Volcano plot of differentially expressed genes among CD8+ T cells comparing ICI-AIN and ICI-ATN, **C.** Gene module score for “Oxidative Phosphorylation” at the niche level. ***: *P* < 0.001 in Mann-Whitney U test, **D.** Pearson correlation between gene module scores for “Oxidative Phosphorylation” and “IFNg Response” in proximal tubule niches from ICI-AIN patients, **E.** Feature plot UMAP of the “IFNg Response” module score mapped on immune niches. Red dashed lines indicate the “proinflammatory myeloid cell” niches. Smaller feature plots show gene expression levels of *MMP9* and *IDO1*, marker genes of proinflammatory myeloid cells, **F.** Violin plot of the “IFNg Response” module score shown for different immune niches, **G.** Likelihood of directly neighboring a fibrosis niche shown for different immune niches. Each dot represents one sample, the dash shows the mean per condition, **H.** Violin plot of the “Fibrotic Niche” module score shown for different immune niches.

As identified by gene set enrichment analysis (**Fig. 4B-D**), gene expression relating to oxidative phosphorylation was reduced in ICI-AIN samples. Using the spatial niche data, we found this reduction localized mostly to the niches related to parts of the kidney tubule (**Fig. 5C**). Tubular epithelial cells rely heavily on oxidative phosphorylation.^29^ IFNγ-signaling within the proximal tubule niches was strongly correlated with a gene signature for oxidative phosphorylation (**Fig. 5D**).

Another hallmark effect of IFNγ is the polarization of myeloid cells into a proinflammatory, tissue-aggressive state, as we observed in the “inflammatory myeloid” cluster (**Fig. 3A-B**). ^30^ Within this cluster, we found the expression of genes typical for a proinflammatory myeloid polarisation (such as *MMP9*, *IDO1*), as well as the highest gene expression signature of an IFNγ response (**Fig. 5E-F**). In summary, CD8^+^ T cell-derived IFNγ was associated with reduced oxidative phosphorylation in tubular cells and a proinflammatory signature in inflammatory myeloid cells.

### Tissue Resident Macrophage Niches Associate with Fibrosis

Finally, we investigated which immune cell niches were associated with fibrosis in ICI-AIN. To this end, we calculated the probability of neighbouring an area of fibrosis for each subtype of immune niche, normalized for the relative abundance of immune niches. The CD8+ infiltrated and tissue-resident myeloid cell niches in ICI-AIN were most likely to neighbor fibrotic niches compared to ICI-ATN group (**Fig. 5G**). Additionally, we defined a fibrotic niche transcriptional signature as the top 100 differentially expressed genes for the fibrosis niches and calculated a signature score for each immune subniche. Consistent with the neighborhood analysis, the tissue-resident myeloid niches had the highest fibrosis signature score (**Fig. 5H**), highlighting their role in a fibrotic tissue response.

## DISCUSSION

Here, we describe the landscape of immune infiltration and fibrosis in the kidneys of patients with ICI-AIN versus ICI-ATN. Using high-resolution spatial transcriptomic analysis, we elucidated key features of ICI-AIN, including a strong activation of IFNγ/STAT1 signaling, a reduction in oxidative phosphorylation in tubular cells, and a heterogeneous immune cell infiltration.

Based on the Tessera algorithm^19^ for the identification of niches with similar transcriptional profiles, we were able to identify biologically meaningful, functional units, thereby circumventing the problems associated with cell segmentation. Some niches corresponded to histoanatomical parts of the kidney like the glomeruli or specific segments of the tubule, allowing us to compare the gene expression within specific kidney regions. Other niche types corresponded to pathological changes including immune cell infiltration and fibrosis. Clustering-based analyses enabled us to investigate the heterogeneity of the immune cell infiltrate in detail, highlighting how TLS-like structures, myeloid cell-dominant inflammation, and fibrotic processes happen in parallel within the ICI-AIN damaged kidney. Analyzing the spatial expression of chemokine/receptor pairs enabled us to understand the mechanisms that determine the architecture of these meta-structures.

Our analyses highlight the central role of CD8^+^ T cells in the pathogenesis of ICI-AIN (**Fig. 6**). T cell infiltration appeared to play an important role in all the immune cell niches we identified, and T cell-related genes were among the most abundant in all niches. These T cells, expressing markers of repeated activation like PD1, CTLA4 and LAG3, seemed to coordinate the recruitment and activation of other immune cells by the expression of CXCL13 (attracting B and T cells and forming TLS-like structures) and IFNγ (inducing an inflammatory myeloid phenotype). These inflammatory myeloid cells, in turn, may have induced tissue damage by producing metallo-proteases like *MMP9* and by recruiting additional CXCR3-expressing inflammatory cells producing CXCL9, CXCL10 and CXCL11.

**Figure 6.**
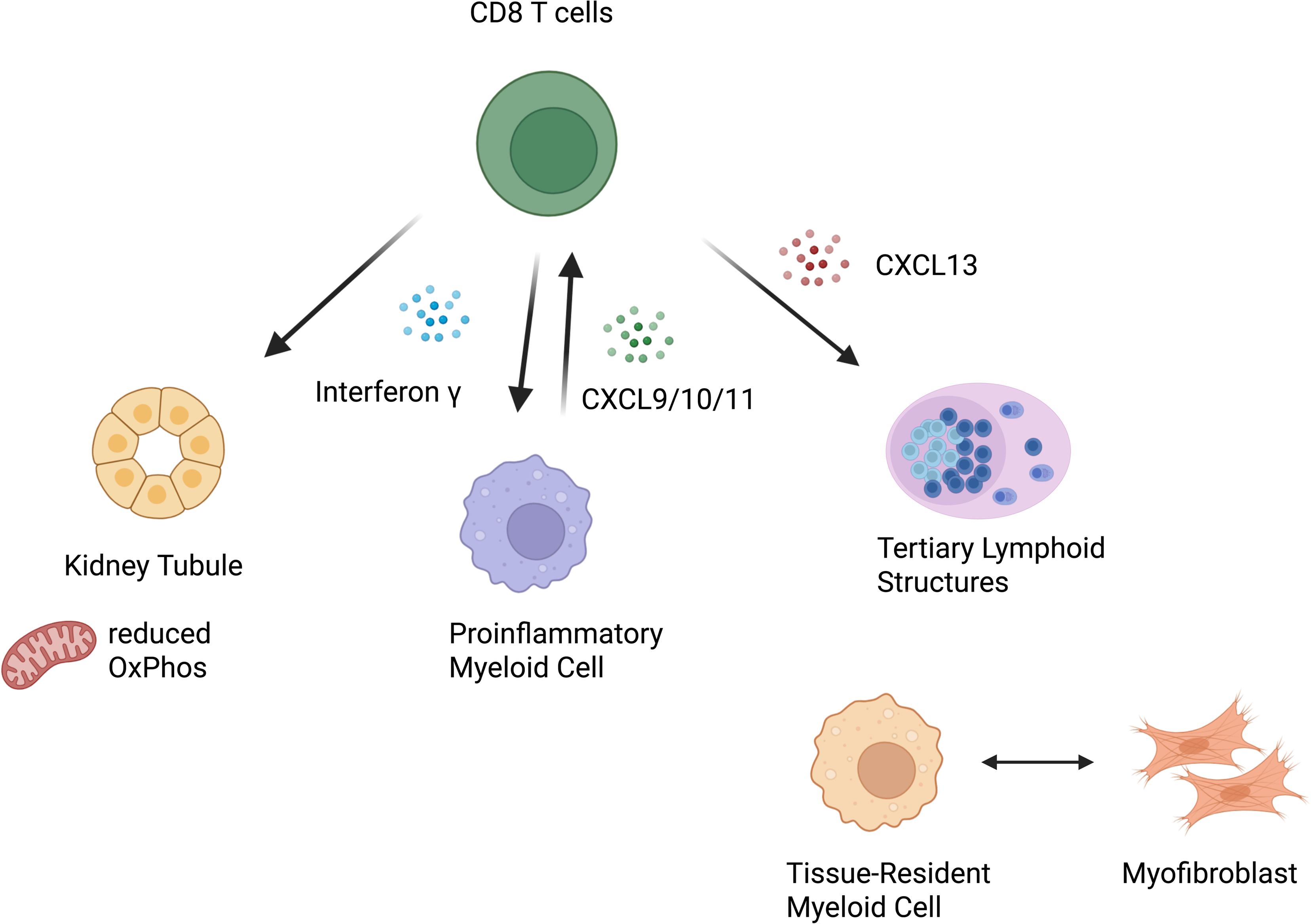
Schematic of the inferred pathophysiology in ICI-AIN with a focus on the role of CD8^+^ T cells: CD8^+^ T cells with transcriptional markers of tissue-residency and repeated stimulation are the main producers of IFNγ. IFNγ reduces oxidative phosphorylation in kidney tubules and induces a proinflammatory program in myeloid cells. These proinflammatory myeloid cells lead to tissue damage and also produce the chemokines CXCL9, CXCL10 and CXCL11, attracting additional CXCR3^+^ lymphocytes, including additional CD8^+^ T cells. CD8^+^ T cells also produce CXCL13, a chemokine that attracts B cells and induces the formation of tertiary lymphoid structures in the kidney.

Notably, CD8^+^ T cells in ICI-AIN recapitulated features associated with T cells of the tumor microenvironment with transcriptional features of exhaustion and instigation of TLS structures.^31–33^ While T cells in ICI-ATN samples showed a transcriptional signature typical of short-lived effector cells (expressing *KLRG1* and *CX3CR1*), the CD8 T cells in ICI-AIN expressed more Hobit (*ZNF683*), CD103 (*ITGAE*) and CXCR6, markers of tissue-resident memory cells.^34^ Interestingly, a CD8^+^ T cell subset with high expression of Hobit was identified in the peripheral blood of a patient with kidney allograft rejection after ICI treatment.^35^ A similar signature of *ITGAE*^+^ tissue-resident T cells was recently reported in ICI-related colitis.^11^ It is therefore plausible that CD8^+^ tissue-resident memory cells, disinhibited by ICI treatment, are the primary cell type that initiate the inflammatory cascade observed in ICI-AIN.

Our analyses also implicated niches with a high density of a tissue-resident transcription profile in the development of kidney fibrosis. This is consistent with recent reports highlighting the role of a similar population in fibrotic damage in lupus nephritis.^36,37^ While we reported the spatial association of these niches with fibrotic area, future research is needed to determine if these cells mediate fibrosis or protect against further damage.

The immunological changes we observed may inform the selection of novel treatment targets in ICI-AIN. Currently, ICI-AIN is treated by holding or discontinuing ICIs and administering glucocorticoids, which may compromise anti-tumor immune response.^38,39^ Drugs targeting IFNγ and its receptors, and especially drugs such as baricitinib and ruloxitinib—which target the downstream signaling of JAK/STAT—may thus have greater utility in treating ICI-AIN.

Though this is the most comprehensive spatial data of ICI-AKI to date, there were limitations. Kidney biopsies from patients with ICI-AIN show great heterogeneity, and the small number of samples analyzed, along with lack of inclusion of “healthy” controls, may have limited our ability to explore key differences. Additionally, methods for cellular segmentation in spatial transcriptomic and proteomic analyses are still an area of active investigation and current spatial transcriptomic approaches result in relatively sparse data. For example, the emphasis on CD8^+^ T cells in our data could, at least in part, be explained by the higher transcript density of *CD8A* compared to *CD4*. While we used a niche-based approach to mitigate these limitations, orthogonal approaches like single-cell transcriptomics in ICI-AIN will be needed to phenotype infiltrating immune cells at higher resolution.

In summary, we used spatial transcriptomics to show that IFN-γ-producing CD8^+^ T cells may be implicated in the pathogenesis of ICI-AIN, and that CD8^+^ T cell-derived IFN-γ likely induce a proinflammatory state in myeloid cells, with increased tissue production of CXCL9, 10, and 11. Future studies with a larger number of samples will be needed to validate our findings and investigate the association of specific immune niche subsets with disease severity/refractoriness, treatment response, and likelihood of kidney recovery.

## Acknowledgments

The authors would like to thank patients and their families who donated to cancer research as part of the Jimmy Fund Walk, as money raised from the walk helped support this work. We would also like to thank the Dana-Farber/Harvard Cancer Center in Boston, MA, for the use of the Specialized Histopathology Core which provided histology and immunohistochemistry service. Dana-Farber/Harvard Cancer Center is supported in part by an NCI Cancer Center Support Grant # NIH 5 P30 CA06516. We acknowledge Ilya Korsunsky and his lab members for assistance with manuscript writing.

## Author Contributions

Designing research studies: SG, KW

Conducting experiments: GC, SW, SIS

Acquiring samples and data: SG, KW, FA, AC, RSR, SAP, RBC, AW, SIS, KM, TB, ACV, NRL, US, KR, ES, DEL, DGM, MES, DR

Analyzing data: QQ, LO, SW, MT, XZ

Writing the manuscript: SG, KW, QQ, LO

All authors critically revised and approved the final version.

## Data Sharing Statement

All data processing and analysis codes are publicly available at https://github.com/qinqian/kidney_autoimmune.

De-identified Xenium data may be available with the data use agreements.

