## Supplemental Information for "Spatial Transcriptomics Identify T Cell-Driven Mechanisms of Kidney Damage in Immune Checkpoint Inhibitor-Associated Acute Interstitial Nephritis"

***Suppl. Figure 7. KEGG and Hallmark Pathway Enrichment Analysis for the Glomerulus, Thick Ascending Limb, and Distal Convoluted Tubule Niches in ICI-AIN and ICI-ATN*** .............................. ***13***

***Supplemental References*** ........................................................................................................... ***14***

### SUPPLEMENTAL METHODS

#### Cell Segmentation

To improve the cell segmentation for potentially missing cells, we ensembled the segmentation results from CellPose[1] and Baysor (v0.7).[2] CellPose-based segmentation was generated using DAPI staining images and provided to Baysor with a set prior confidence of 1.0. The whole slide was divided into small patches for building the model in a plausible memory. Segmentations were then merged and resolved to match the original slide, and output genes by cell counts matrix with spatial coordinates. The overall pipeline was implemented in SOPA (v2.0.6),[3] and driven by Snakemake.[4] We further checked the Xenium cells' quality by cell numbers, transcript numbers in total and per cell, and excluded cells with fewer than 20 feature counts and 20 transcripts counts.

#### Cell Annotation

We used the kidney single-cell RNA-sequencing (scRNA-seq) atlas from the Kidney Precision Medicine Project (KPMP) as a reference for cellular annotation.[5–7] We then normalized gene expression from the combined single-cell and spatial transcriptomic data with an average scaling factor of median counts in each. Principal components analysis with 50 components was performed on the scaled expression of top variable genes. Using the 50 components, we integrated the scRNA-seq from the KPMP with spatial transcriptomic datasets, and corrected for batch effects by assigning batch variables to techniques and samples in Harmony with default parameters.[8] The labels were transferred from the KPMP single cell atlas to our generated Xenium data by assigning the major cell types with the labels based on their 10 nearest neighbors. Nephrologists, renal pathologists, and immunologists verified and refined the cell subtypes.

#### Marker Identification

With the existing clustering of spatial niches or cells, differentially expressed genes (DEGs) were further identified by comparing a given cluster with the remaining clusters using Wilcoxon signed-rank test through the presto package (https://github.com/immunogenomics/presto). Markers were then used for curating cell types or annotating niche types *de novo*. The same statistical test was also applied when comparing the curated spatial niches between the ICI-AIN and ICI-ATN groups.

**Spatial Niche Identification and Annotation**

We identified spatial niches using the Tessera package.[9] We limited the "tile" size to 5 to 20 micrometers to aggregate the spatially-adjacent cells with similar. The aggregated cells were normalized and dimensionally reduced using the same workflow as in **Section Cell Annotation**. Niches were further classified based on consensus annotation by two nephrologists, and validated using the aligned H&E images. By aggregating the cells into niches, we reduced the data sparsity bias in the Xenium platform, and we reproduced the expected kidney biology in spatial niches. Subsetting the immune niches, we performed re-harmonization and unbiased clustering. Based on prior knowledge and differentially expressed genes across clusters, we further annotated each cluster.

**Signaling Cross-Talk Analysis**

We analyzed spatial niche interactions and crosstalk through customized scripts with spatula[10] and CellChat.[11] First, we applied spatula to define the neighborhood of spatial niches based on the centroids, and we quantified the interaction and normalized by the neighbors number in a given distance (30 micrometer). A permutation test with 100 permutations across space was made to compute the Z-score, p-value, and false discovery rate (FDR), with an interaction FDR $< 0.05$ considered significant. Then, CellChat was applied with the CXCL and CCL signaling pathways in the CellChatDB human database to identify enrichment of the ligand receptor pairs in the spatial transcriptomics data.

**Gene Set Enrichment Analysis**

Gene set pathway enrichment analysis was performed using the fgsea package[12] with the pre-compiled KEGG pathway and Hallmarks gene sets in Molecular Signature Database (MSigDB) R package msigdbr.[13] The top enriched pathways were filtered by adjusted p-value of 0.05 and ranked by the adjusted p-value. The pathway-specific scores were summarized using UCell[14] and projected onto spatial niches.

**Differential Cell Abundance Analysis**

We applied differential cell abundance analysis in two ways: covarying neighborhood and linear regression analysis to identify the compositional differences of cells and niches between ICI-AIN and ICI-ATN. First, we used the covarying neighborhood analysis (cna) method[15] for single-cell level differential abundance test in a cluster-free approach. Second, we applied the sccomp[16] to the cluster-level differential abundance test.

**Visualizations**

The R packages dittoSeq (v1.20.0)[17] and tidyplots (v0.3.1)[18] were used to produce visualizations for the paper. Figure 1A and Figure 6 were created in BioRender.

### Suppl. Table 1. Characteristics of Xenium Cohort

| Pt | Age/ Sex | Comorbidities | eGFR[a] | Cancer Type | ICI | SCr BL/Peak/ Nadir[b] (mg/dL) | Time to AKI (weeks)[c] | Bx | Proteinuria (dipstick/ UPCR) | LE/Blood (dipstick) | Extra-renal irAE[d] | Concomitant NSAID or PPI use[e] | Concomitant Chemotherapy[f] |
|---|---|---|---|---|---|---|---|---|---|---|---|---|---|
| 1 | 68/F | HTN | 53 | GU | Pembro | 1.1/2.8/1.2 | 38 | **AIN** | Neg/0.18 | Neg/Neg | None | None | No |
| 2 | 66/F | DM/COPD | 97 | Lung | Pembro | 0.6/3.8/1.9 | 30 | **ATN** | NA | NA | None | PPI | Carboplatin/ pemetrexed |
| 3 | 83/F | HTN/HLD | 36 | GU | Pembro | 1.4/2.7/1.0 | 49 | **AIN** | NA | NA | None | None | No |
| 4 | 79/M | None | 59 | Head and Neck | Pembro | 1.3/2.0/1.3 | 9 | **ATN** | 1+/0.18 | Neg/2+ | None | NSAIDs | Lupron/ Letrozole/ Ribociclib |
| 5 | 68/F | HTN/HLD/DM | 73 | Uterine | Pembro | 0.9/2.2/1.1 | 6 | **AIN** | 1+/0.49 | 1+/Neg | Rash | NSAIDs | Lenvatinib |
| 6 | 55/F | None | 92 | Melanoma | Atezo | 0.8/2.2/1.0 | 3 | **AIN** | Neg/0.74 | 1+/3+ | None | None | Bevacizumab |
| 7 | 75/M | HLD | 60 | NSCLC | Pembro | 1.3/3.1/1.7 | 20 | **ATN** | NA | NA | Colitis | None | Cisplatin/ pemetrexed |
| 8 | 76/M | HLD | 63 | RCC | Nivo/ Relatlimab | 1.2/1.9/1.4 | 39 | **ATN** | Neg/0.08 | 1+/Neg | None | None | Zanzalintinib |

Abbreviations: AIN, acute interstitial nephritis; AKI, acute kidney injury; Atezo, atezolizumab; ATN, acute tubular necrosis; BL, baseline; bx, biopsy; COPD, chronic obstructive pulmonary disease; DM, diabetes mellitus; GU, genitourinary; HLD, hyperlipidemia; HTN, hypertension; ICI, immune checkpoint inhibitor; irAE, immune-related adverse events; LE, leukocyte esterase; Neg, negative; Nivo, nivolumab; NSAID, nonsteroidal anti-inflammatory drugs; NSCLC, non-small cell lung cancer; Pembro, pembrolizumab; PPI, proton pump inhibitor; RCC, renal cell carcinoma; SCr, serum creatinine; UPCR, urine protein to creatinine ratio

[a]Baseline eGFR was defined based on the SCr value closest and prior to ICI initiation, and was calculated based on Chronic Kidney Disease-Epidemiology Collaboration equation without race.

[b]Nadir refers to the lowest serum creatinine value achieved within 90 days following AKI onset.

[c]Weeks from ICI initiation to AKI onset

[d]Extra-renal irAE refers to those occurring in the 14 days preceding or following onset of AKI

[e]Concomitant NSAID or PPIs refers to use in the 14 days preceding or following onset of AKI

[f]Concomitant chemotherapy refers to receipt in the 30 days preceding onset of AKI

All patients with ICI-AIN received glucocorticoids

### Suppl. Figure 1. Single Cell Analysis of Spatial Transcriptomic Data in ICI-AIN and ICI-ATN

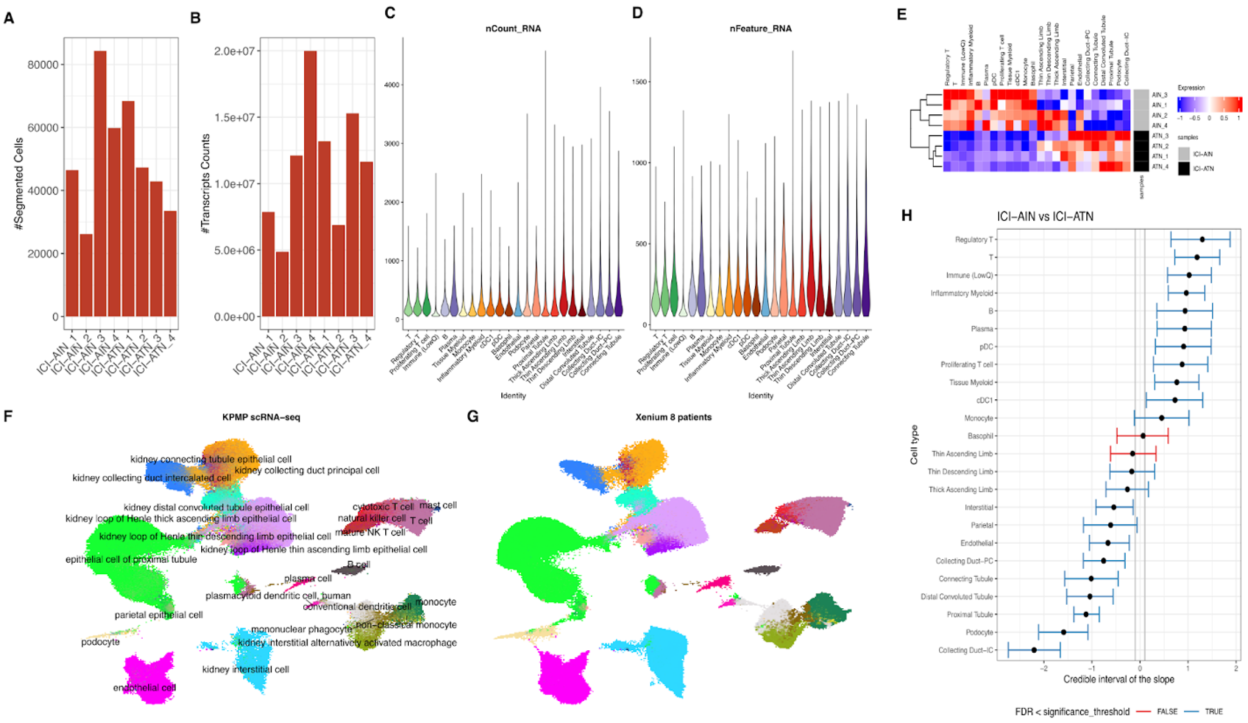

**A.** Bar chat shows the segmented cells per sample using CellPose plus Baysor**, B.** The bar chart shows the transcripts counts per sample, **C.** The violin plot showing the number of transcripts counts per cell type**, D.** The violin plot shows the number of feature counts per cell type**, E.** Heatmap showing the scaled cell type ratio across the samples, **F-G.** KPMP single cell RNA-seq integration with spatial transcriptomic samples, **H.** sccomp analysis by comparing ICI-AIN vs ICI-ATN groups.

### Suppl. Figure 2. Representative Biomarkers for the Major Cell Types in Fig. 1

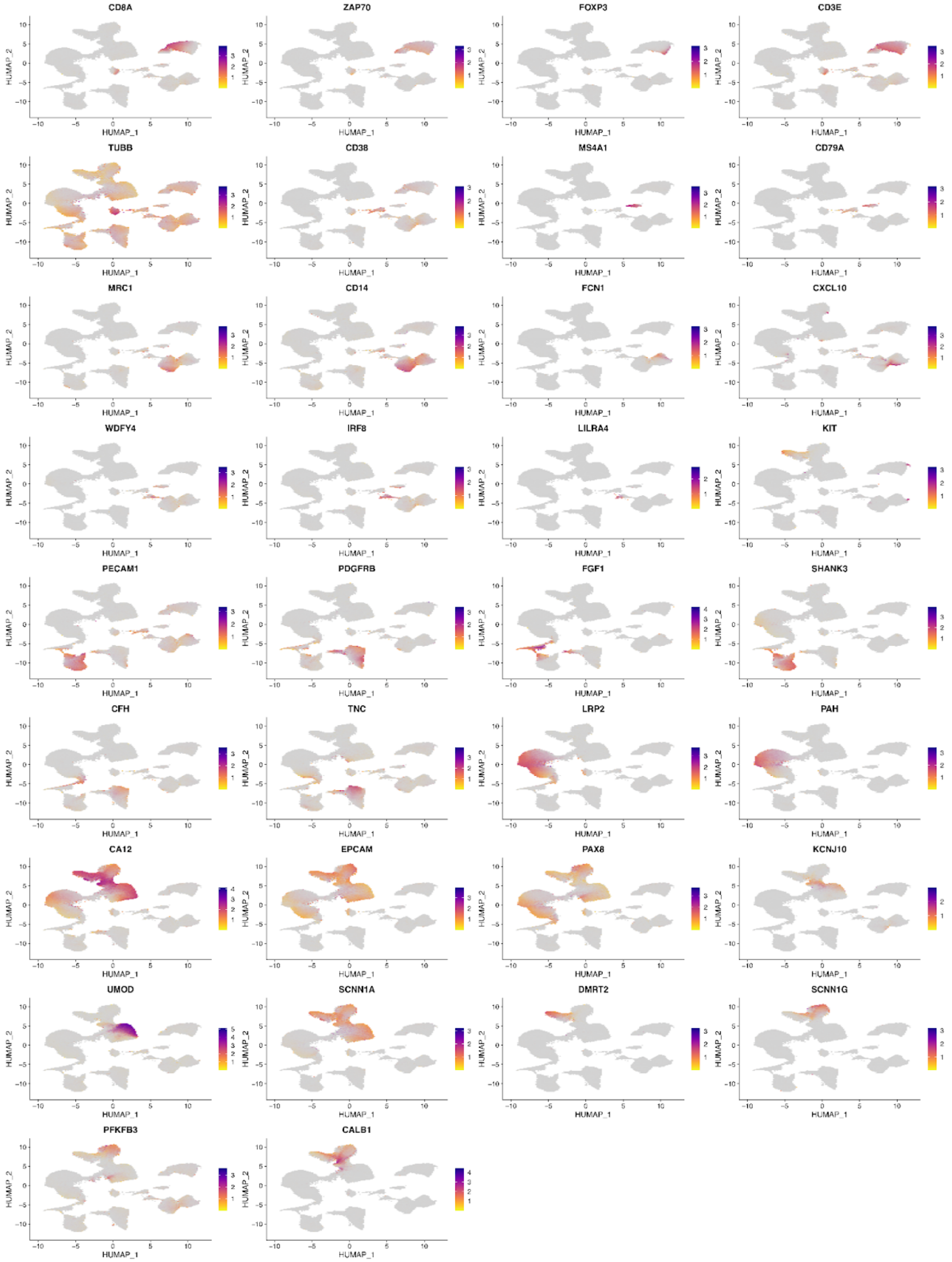

**Suppl. Figure 3. Validation of Tessera-Defined Niches Using H&E. Zoom-in Plot of Tessera Niches, Niche-Specific Marker Expression, and Aligned H&E Imaging.**

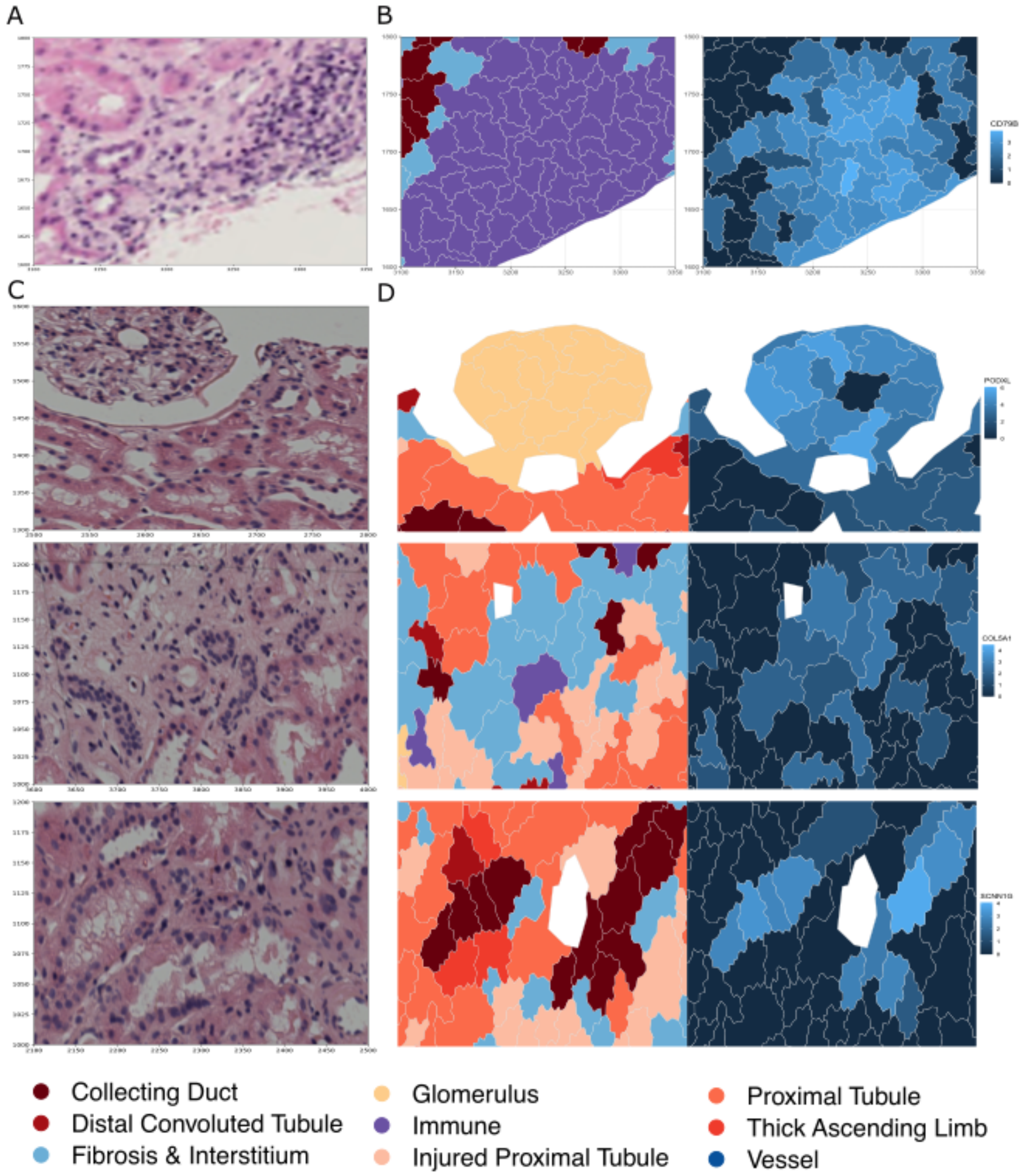

A. ICI-AIN patient’s H&E imaging of the immune niche
B. ICI-AIN patient’s Xenium-based immune niche together with the CD79B expression distribution in niche, color legend is the same as in the Fig. 2
C. ICI-ATN patient’s zoom-in example patches of H&E imaging, from top to bottom panels are examples of glomerulus niche, Fibrosis & Interstitium niche, and collecting duct niche
D. The similar zoom-in plot for the Xenium niche analysis in the ICI-ATN patient in C

### Suppl. Figure 4. CellChat Analysis of Niche-Based Crosstalk.

#### CellChat crosstalk analysis

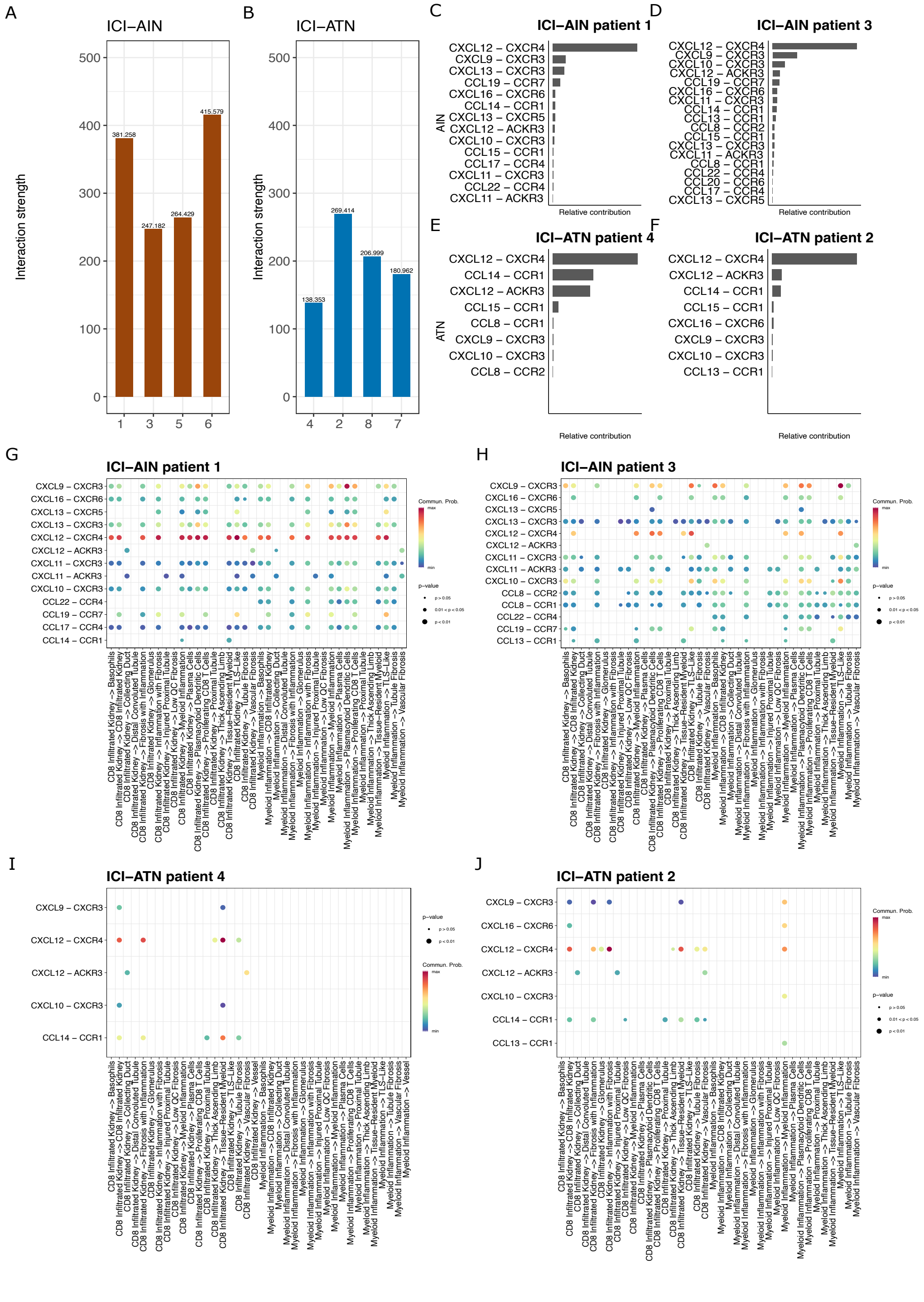

**A-B.** Overall ligand-receptor interaction frequencies in ICI-AIN and ICI-ATN. **C-D.** ICI-AIN CXCL and CCL ligand-receptor pairs' contribution to niche-based crosstalk. **E-F.** the same analysis in the ICI-ATN group. **G-H**. ICI-AIN analysis of CXCL and CCL ligand-receptor pairs interaction probability in different niche pairs starting from CD8 Infiltrated Kidney or Myeloid Inflammation niches**, I-J.** the same analysis of representative samples in ICI-ATN

### Suppl. Figure 5. Comparison of Olink Proteomics and Xenium Data

**A**

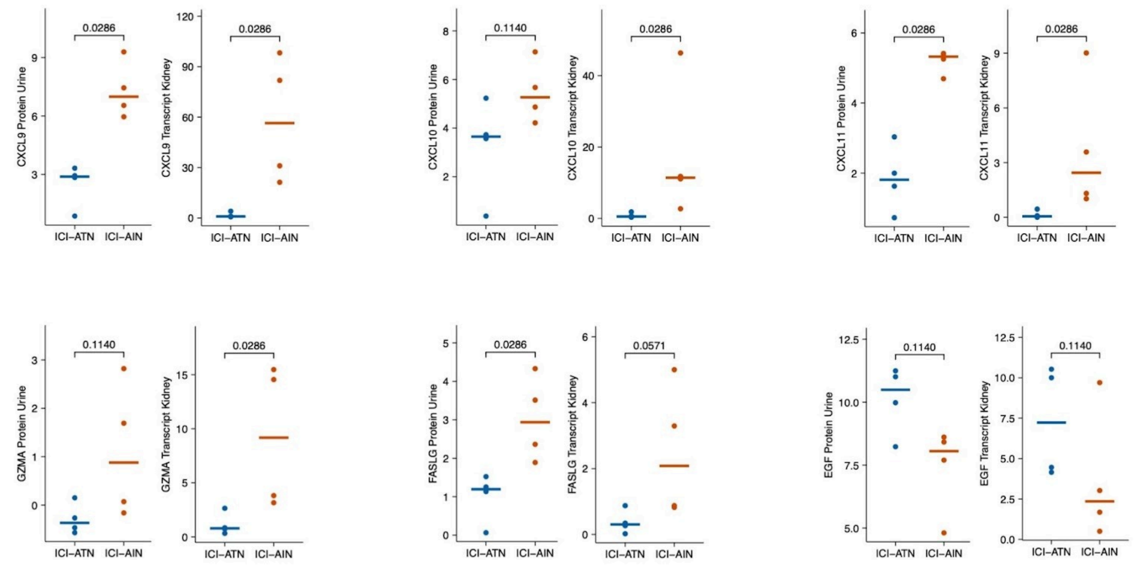

Analysis of protein concentration in the urine (as measured in Olink assay, units are Normalised Protein Expression – NPX) and aggregate gene expression per area in the Xenium kidney spatial transcriptomics. *P* values calculated using Mann-Whitney U test.

**Suppl. Figure 6. KEGG and Hallmark Pathway Enrichment Analysis for 6 Spatial Niches in ICI-AIN vs. ICI-ATN.**

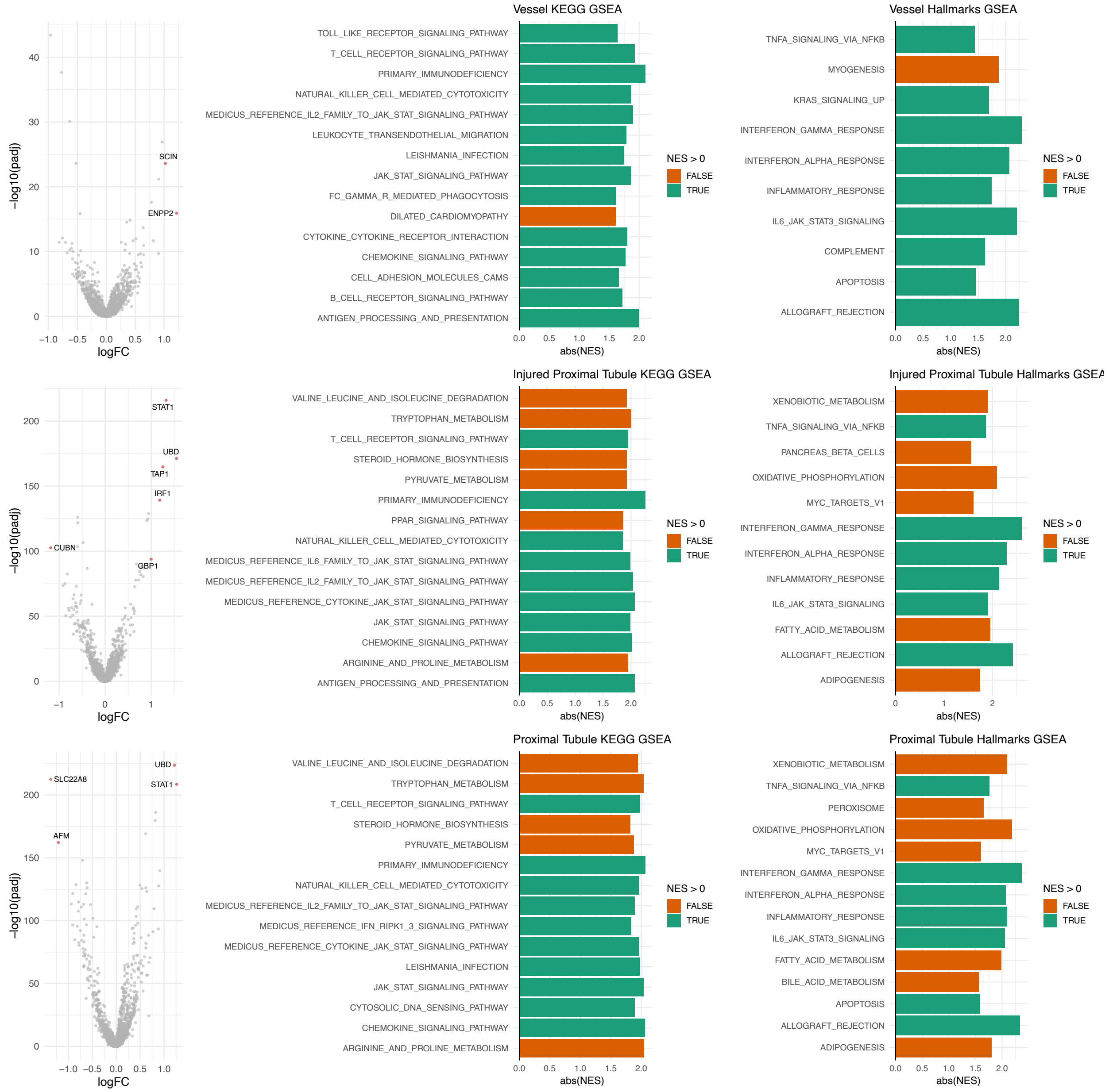

**Suppl. Figure 7. KEGG and Hallmark Pathway Enrichment Analysis for the Glomerulus, Thick Ascending Limb, and Distal Convoluted Tubule Niches in ICI-AIN and ICI-ATN**

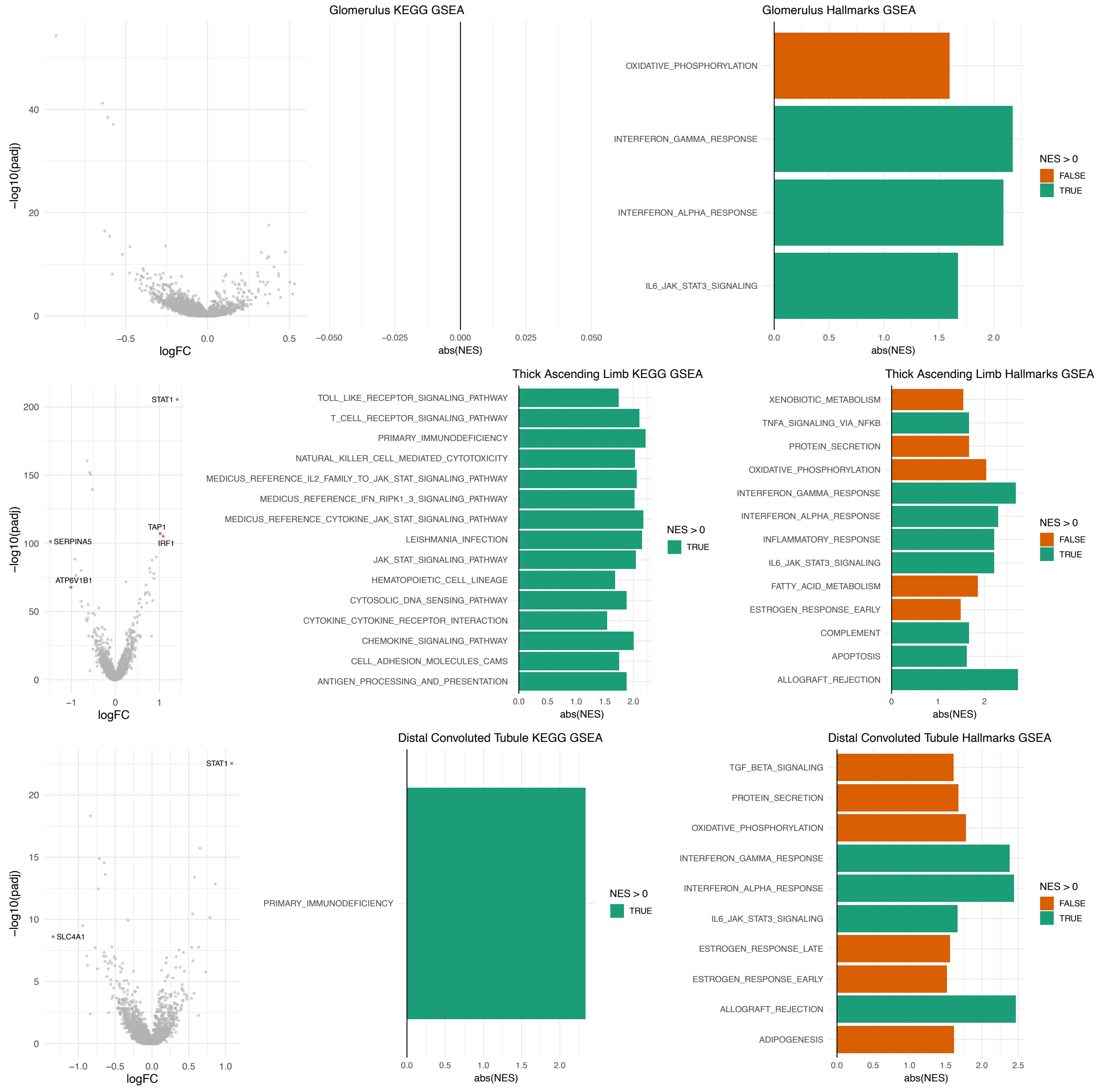
